# Septin7 is essential in early hematopoiesis, but redundant at later stages

**DOI:** 10.1101/2025.04.22.649981

**Authors:** Natalia Ronkina, Nikita A. Verheyden, Prerna Gambhir, Kathrin Laaß, Tatiana Yakovleva, Anneke Dörrie, Megha Abbey, Manoj B. Menon, Anton Selich, Melanie Galla, Michael Rothe, Axel Schambach, Andreas Krueger, Alexey Kotlyarov, Matthias Gaestel

## Abstract

The unique cytoskeletal protein Septin7 is generally considered to be required for cytokinesis in yeast and mammals. Whole body genetic ablation of *Septin7* in mice is embryonic lethal. *Septin7-*deficient fibroblasts and HeLa cells are defective in cytokinesis and undergo obligate multinucleation. Surprisingly, lymphocyte- and myeloid-specific deletion of *Septin7* in mice did not result in any detectable abnormalities in blood lineage development, suggesting that Septin7 is dispensable for steady-state hematopoiesis. In contrast, *Septin7*-deficient hematopoietic stem cells failed to engraft and establish donor chimerism following transplantation, indicating that Septin7 is essential for hematopoietic stem cell function under stress conditions. To reconcile these contradictory findings, we analyzed the effects of *Septin7* deletion in hematopoietic cells from mice with either pan-hematopoietic or lymphoid lineage-specific *Septin7* deletion. Our results demonstrate that Cre-induced deletion of the floxed *Septin7* allele is inefficient during the early stages of hematopoiesis, suggesting strong selection pressure against Septin7 deficiency at this stage. In contrast, deletion of *Septin7* at the common lymphoid progenitor stage, as well as in Hoxb8-immortalized hematopoietic progenitors, is efficient and does not result in any noticeable defects in cell division. Taken together, our findings indicate that Septin7 is essential during early hematopoiesis but becomes dispensable at later stages.

## Introduction

Cytokinesis in yeast and animals depends on the contraction of the cell membrane at an equatorial furrow. This contraction is powered by a ring-like structure composed of actin, myosin, and several cytoskeleton-associated proteins. Effective cytokinesis is achieved by a comprehensive coordination of cell-cycle control, spatial organization, rearrangement of the cytoskeleton, generation of mechanical forces, and the trafficking of cellular membranes (Hartwell et al. 1970; Marquardt et al. 2021; Addi et al. 2018; Fededa and Gerlich 2012; Green et al. 2012; Karasmanis et al. 2019). Initially, identified as essential genes for cytokinesis in budding yeast, septins have been found to interact with the mitotic spindle, the contractile ring, and the midbody, playing a crucial role in the cytokinesis process of metazoan cells (Fededa and Gerlich 2012; Green et al. 2012; Karasmanis et al. 2019; Hartwell et al. 1970). Mammalian cells express 13 different septins, which can be divided into four groups based on sequence similarity. Three of these septin groups are represented by multiple members, but the fourth group consists only of the single member Septin7 (SEPT7). Members of the four groups assemble into septin octamers, which polymerize to linear, nonpolar polymers. In this structure, septins of the same group can substitute for each other, but SEPT7, as the only member of the fourth group, cannot be substituted by other septins. Therefore, the absence of SEPT7 prevents octamer formation, destabilizes other core septins and causes loss of functional septin filaments. In the past, we generated and characterized *Sept7* conditional knockout mice (Menon et al. 2014). *Sept7*^Δ/Δ^ embryos were detected in utero only up to embryonic day 6.5 (E6.5)-E7.0 but not later (Menon et al. 2014). SEPT7-deficient fibroblasts and HeLa cells display cytokinetic failure and undergo obligate multinucleation indicating a crucial role of SEPT7 in cell division (Menon et al. 2014; Kremer et al. 2005). Unexpectedly lymphocyte specific targeting of *Sept7* (Menon et al. 2014) and myeloid specific targeting (Menon et al. 2022) demonstrated no overt abnormality in the development of the blood cell lineages, leading us to the conclusion that SEPT7 is not required for hematopoiesis. However, later study of *Vav1*-iCre-driven deletion of *Sept7* in hematopoietic stem and progenitor cells (HSPCs), which sit at the apex of the hematopoietic system to form the major blood lineages, demonstrated dramatically reduced engraftment potential along with characteristics of aged hematopoietic stem cells (HSCs) and decreased lymphoid-primed multipotent progenitors in the bone marrow (Kandi et al. 2021). Here, we analyze the role of SEPT7 in hematopoietic cells at different stages of development. Our results reveal a stage-specific role of SEPT7 in hematopoiesis, demonstrating that while SEPT7 is crucial for early hematopoiesis, it becomes dispensable in later stages.

## Results

### Hematopoietic cells escape *Vav1*-iCre*-*mediated recombination of the *Sept7* ^flox/flox^ allele

To establish the role of SEPT7 in hematopoiesis, we aimed to conditionally delete *Sept7* in mice by taking advantage of hematopoiesis-restricted Cre recombinase (Cre) trans-gene expression in *Vav1*-iCre mice (de Boer et al. 2003). We crossed loxP-flanked *Sept7* (*Sept7*^flox/flox^ or *Sept7*^wt/flox^) mice with *Vav1-iCre* mice. *Vav1-iCre* expresses active Cre in hematopoietic stem and progenitor cells and all hematopoietic descendants(de Boer et al. 2003). Genotyping analysis of splenocytes, thymocytes and bone marrow from *Vav1-iCre*::*Sept7*^wt/flox^ and *Vav1-iCre*::*Sept7*^flox/flox^ mice demonstrated incomplete excision of *Sept7* floxed alleles from *Vav1-iCre*::*Sept7*^flox/flox^ mice (Figure 1A). In contrast, no traces of *Sept7* floxed allele could be detected in *Vav1-iCre*::*Sept7*^wt/flox^ cells, indicating highly effective Cre-mediated excision of *Sept7* floxed allele in the cells, carrying one wild type allele of *Sept7* (Figure 1A). Western blot analysis of SEPT7 protein expression revealed clearly detectable SEPT7 protein levels in splenocytes, thymocytes and bone marrow isolated from *Vav1-iCre*::*Sept7*^flox/flox^ mice (Figure 1B). Intracellular SEPT7 protein expression assessed by flow cytometry demonstrated, that most of splenocytes, thymocytes and bone marrow cells from *Vav1-iCre*::*Sept7*^flox/flox^ mice express SEPT7 (Figure 1C). Some portion of SEPT7 deficient cells could be observed in thymocytes of both *Sept7*^flox/flox^ mice and in splenocytes from one of the two *Sept7*^flox/flox^ mice (Figure 1C). We therefore focused our analysis on intrathymic T-cell development. We hypothesized that periodic strong proliferative bursts followed by cell-cycle inactivity during somatic recombination of antigen-receptor genes might reveal proliferation-driven selection pressure. Overall development of T cells from *Vav1-iCre*::*Sept7*^flox/flox^ mice was unimpaired (Figure S1). We observed similar proportions of double-negative (DN), double-positive (DP) and single-positive (SP) thymocyte populations and similar frequencies of CD4^+^ and CD8^+^ T cells as well as B cells within spleen of *Vav1-iCre*::*Sept7*^wt/flox^ and *Vav1-iCre*::*Sept7*^flox/flox^ mice. Analysis of SEPT7 protein in thymocyte subsets covering stages of T-cell receptor gene rearrangement (DN3a, pre-selection DP), proliferative stages (DN3b, DN4), and selection/maturation stages (post-selection DP) revealed biphasic distribution of SEPT7 in cells from *Vav1-iCre*::*Sept7*^flox/flox^ mice. Thus, cells with incomplete deletion of floxed Sept7 alleles, referred to as escapees, are present at each developmental stage. However, we note that SEPT7-negative cells are maintained even through stages of high proliferative activity (DN3b and DN4) (Figure 1D). Considering the complete excision of *Sept7* floxed alleles in *Vav1-iCre*::*Sept7*^wt/flox^ mice, we conclude that hematopoietic cells are strongly driven to evade *Vav1*-iCre-mediated recombination of the *Sept7*^flox/flox^ allele due to intense selection pressure favoring the survival of SEPT7 expressing cells (Figure 1E).

**Figure 1.**
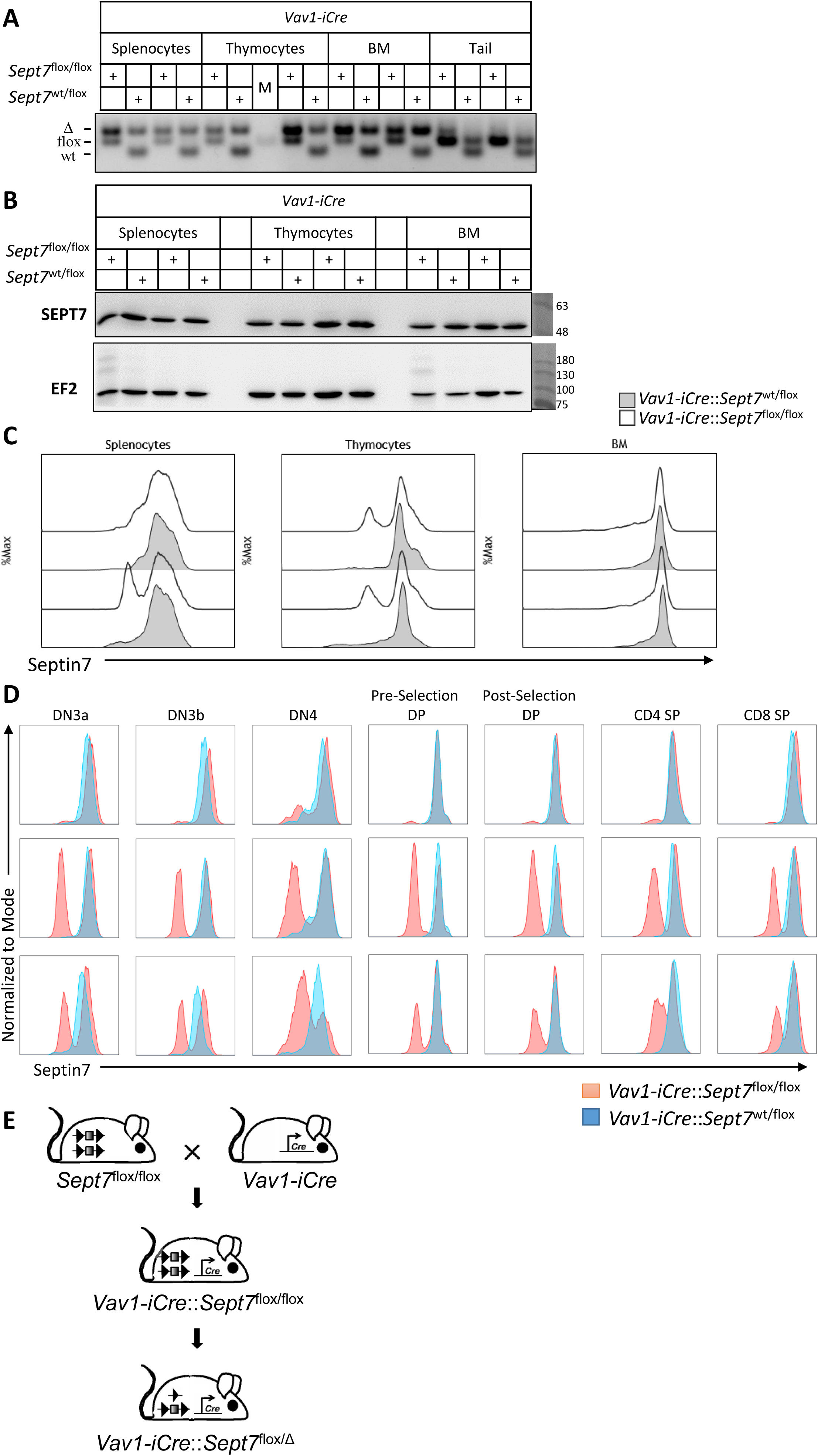
Hematopoietic cells escape *Vav1*-iCre*-*mediated recombination of *Sept7*^flox/flox^ allele. Splenocytes, thymocytes and bone marrow cells (BM) as well as tail biopsy (Tail) were isolated from *Vav1-iCre*::*Sept7*^wt/flox^ and *Vav1-iCre*::*Sept7*^flox/flox^ littermates. Two different mice of each genotype were used. A. Genotyping reveals complete excision of *Sept7* floxed allele in *Vav1-iCre*::*Sept7*^wt/flox^ mice and not efficient excision of *Sept7* floxed allele in *Vav1-iCre*::*Sept7*^flox/flox^ mice. B. Western blot analysis demonstrates expression of SEPT7 in *Vav1-iCre*::*Sept7*^flox/flox^ cells, which are expected to be SEPT7 deficient. C. FACS analysis indicates only minor fraction of SEPT7 deficient thymocytes and splenocytes isolated from *Vav1-iCre*::*Sept7*^flox/flox^ mice: vast majority of the cells escape *Vav1-iCre*-induced recombination and express SEPT7. D. FACS analysis of SEPT7 expression at different stages of T-cell development. The escapees are detected at all stages of T-cell development in *Vav1-iCre*::*Sept7*^flox/flox^ mice. Three mice of each genotype were analyzed. E. Schematic presentation of the experimental results: incomplete excision of *Sept7* floxed allele in hematopoietic cells from *Vav1-iCre*::*Sept7*^flox/flox^ mice.

### SEPT7 is effectively eliminated in T-cells from *hCD2-iCre*::*Sept7*^flox/flox^ mice

To assess, whether SEPT7 negative cells are indeed capable of undergoing T-cell development, we generated mice in which Cre-mediated excision of *Sept7* was restricted to T- and B-cells. We crossed loxP-flanked *Sept7* (*Sept7*^flox/flox^ or *Sept7*^wt/flox^) mice with *hCD2-iCre* mice. *hCD2-iCre* expresses active Cre in immature and mature B and T lymphocytes beginning at the common lymphoid progenitor stage (CLP) (de Boer et al. 2003). Genotyping analysis of splenocytes, thymocytes and T-cells from *hCD2-iCre*::*Sept7*^wt/flox^ and *hCD2-iCre*::*Sept7*^flox/flox^ mice demonstrated effective excision of *Sept7* floxed alleles in T-cells of *hCD2-iCre*::*Sept7*^flox/flox^ mice (Figure 2A). Western blot analysis of SEPT7 protein expression revealed significantly decreased SEPT7 protein levels in splenocytes and thymocytes of *hCD2-iCre*::*Sept7*^flox/flox^ when compared to *hCD2-iCre*::*Sept7*^wt/flox^ mice or to the *Sept7*^flox/flox^ mice without Cre-expression (Figure 2B). Flow cytometric analysis demonstrated that the depletion of SEPT7 at the DN2/DN3a stage was not complete (Figure 2C). In DN3b thymocytes the SEPT7 protein seems to be largely absent and even less SEPT7 could be detected in DN4, although both populations proliferate strongly (Figure 2C). In DP cells, a small proportion of *hCD2-iCre*::*Sept7*^flox/flox^ cells was SEPT7-positive prior to selection and a few CD4 SP and CD8 SP cells may have retained SEPT7 expression as assessed by flow cytometry as well (Figure 2C). There were no differences in T-cell differentiation between *hCD2-iCre*::*Sept7*^flox/flox^ and *hCD2-iCre*::*Sept7*^wt/flox^ in thymus - all populations of thymocytes were comparable in both animals (Figure S2) . Given that upon virtually complete deletion at the DN3a stage of T-cell development, no substantial frequencies of escapees emerged in this experimental model, we conclude that SEPT7 is not essential for proliferation during T-cell development at and after the DN3 stage (Figure 2D). Our observation is consistent with previous studies demonstrating that intrathymic development of SEPT7 deficient T-cells remains largely unaffected (Mujal et al. 2016; Menon et al. 2014).

**Figure 2.**
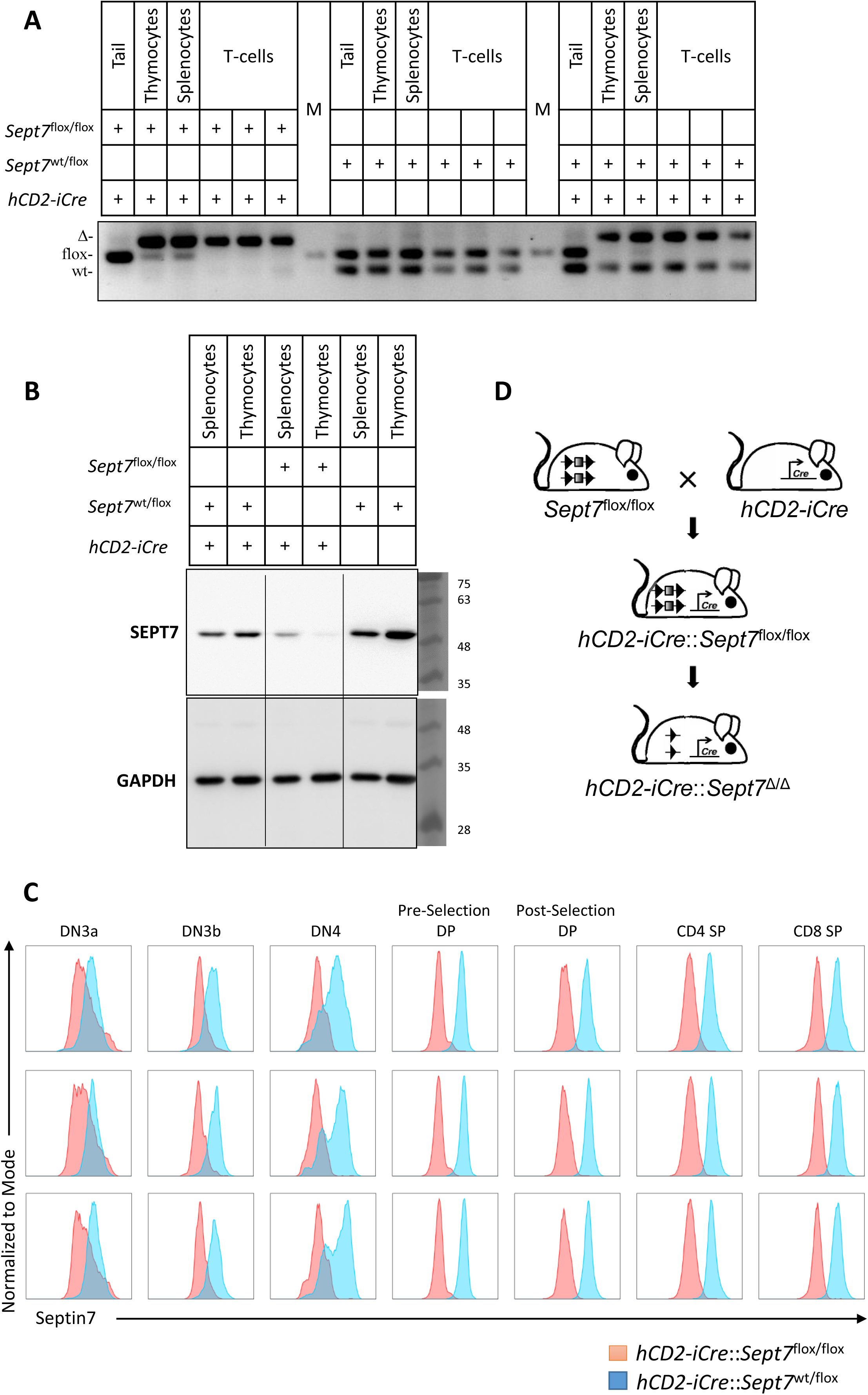
SEPT7 is effectively eliminated in T-cells from *hCD2-iCre*::*Sept7*^flox/flox^ mice. Splenocytes, thymocytes, T-cells and tail biopsy (Tail) were isolated from *hCD2-iCre*::*Sept7* ^wt/flox^ and *hCD2-iCre*::*Sept7* ^flox/flox^ littermates and from *Sept7* ^wt/flox^ mice without *hCD2-driven iCre* expression. A. Genotyping reveals effective excision of *Sept7* floxed allele in T-cells from *hCD2-iCre*::*Sept7* ^wt/flox^ and *hCD2-iCre*::*Sept7* ^flox/flox^ mice. B. Western blot analysis demonstrates decreased expression of SEPT7 in *hCD2-iCre*::*Sept7*^flox/flox^ thymocytes and splenocytes. C. FACS analysis indicates accumulation of SEPT7 deficient cells during T-cell development. Three different mice of each genotype were analyzed. D. Schematic presentation of the experimental results: complete excision of *Sept7* floxed allele in T-cells from *hCD2-iCre*::*Sept7*^flox/flox^ mice

### Hoxb8 immortalized HSPCs from *Vav1-iCre*::*Sept7*^flox/flox^ mice escape Cre-mediated recombination and express SEPT7

To generate a stably growing, homogenous hematopoietic progenitor cell population we immortalized early hematopoietic stem and progenitor cells from *Vav1-iCre*::*Sept7*^flox/flox^ and *Vav1-iCre*::*Sept7*^wt/flox^ mice by gammaretroviral delivery of estrogen–regulated Hoxb8 (Redecke et al. 2013). Monitoring of Cre-mediated recombination in these hematopoietic cells by genotyping revealed that *Vav1-iCre*::*Sept7*^flox/flox^ cells had incomplete recombination of the *Sept7* floxed alleles suggesting the presence of cells that had retained at least one functional *Sept7* allele (Figure 3A). No traces of *Sept7* floxed allele could be detected in *Vav1-iCre*::*Sept7*^wt/flox^ cells, indicating highly effective Cre-mediated excision of the *Sept7* floxed allele in the cells, where cell growth was protected by the presence of wild type allele of *Sept7* (Figure 3A). Western blot analysis of SEPT7 protein expression indicated comparable SEPT7 protein levels in HSPCs of both genotypes (Figure 3B). Flow cytometry analysis of intracellular SEPT7 protein expression revealed that the Hoxb8-immortalized HSPCs derived from *Vav1-iCre*::*Sept7*^flox/flox^ HSPCs express SEPT7 (Figure 3C). Population doubling time of *Sept7*^wt/flox^ and *Sept7*^flox/flox^ cells was comparable (Figure 3D). We also performed limiting cell dilution of *Vav1-iCre*::*Sept7*^flox/flox^ cells to track the fate of individual cells. A total of 16 clones were genotyped after 2-3 weeks of propagation, and the results confirmed that all retained the *Sept7* floxed allele (Figure 3E). Flow cytometry analysis in two random clones (C1 and C3) demonstrates that these clones express SEPT7 (Figure 3F). These results imply a critical role of SEPT7 for stem cell maintenance and this conclusion is strongly supported by our recent study where HSPCs from *Vav1-iCre*::*Sept7*^flox/flox^ mice showed strongly impaired repopulation activity upon transplantation and failed to establish donor chimerism (Kandi et al. 2021).

**Figure 3.**
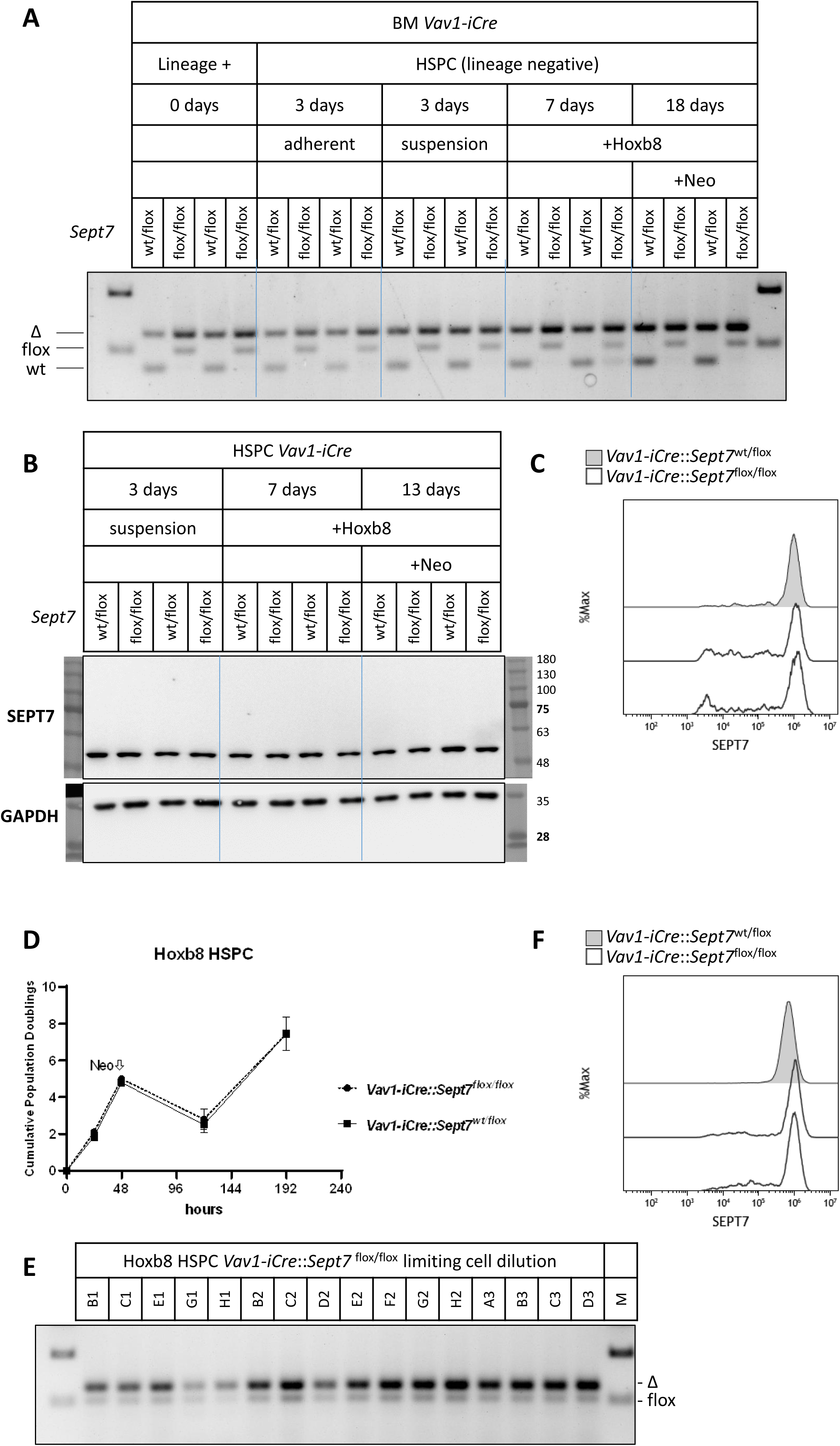
HSPCs escape *Vav1-iCre-*mediated recombination of *Sept7*^flox/flox^ alleles. Lineage-negative bone marrow cells were isolated from *Vav1-iCre*::*Sept7*^wt/flox^ and *Vav1-iCre*::*Sept7*^flox/flox^ littermates. Two different mice of each genotype were used. Cells were stably transduced with Hoxb8 expressing construct that promotes myeloid progenitor proliferation. A. Genotyping reveals complete excision of *Sept7* floxed from *Vav1-iCre*::*Sept7*^wt/flox^ cells and inefficient excision of *Sept7* floxed allele in *Vav1-iCre*::*Sept7*^flox/flox^ cells. B. Western blot analysis demonstrates expression of SEPT7 in *Vav1-iCre*::*Sept7*^flox/flox^ cells, which are expected to be SEPT7 deficient. C. FACS analysis indicates SEPT7 expression in all Hoxb8-immortalized HSPCs derived from *Vav1-iCre*::*Sept7*^flox/flox^ mice. D. Hoxb8-immortalized HSPCs from *Vav1-iCre*::*Sept7*^flox/flox^ mice demonstrate proliferation rate comparable to *Vav1-iCre*::Sep7^wt/flox^ cells. E. Genotyping analysis of single cell colonies derived from Hoxb8-immortalized *Vav1-iCre*::*Sept7*^flox/flox^ HSPCs demonstrates incomplete recombination of *Sept7* floxed allele. F. FACS analysis confirms SEPT7 expression in two random clones (C1 and C3) of Hoxb8-immortalized *Vav1-iCre*::*Sept7*^flox/flox^ HSPCs.

### The *Sept7* floxed allele is effectively protected from Cre-mediated excision in *Vav1-iCre*::*Sept7*^flox/flox^ Hoxb8-immortalized HSPCs

Since the *Sept7* floxed allele is not efficiently recombined in HSPCs from *Vav1-iCre*::*Sept7*^flox/flox^ mice, possibly due to selection pressure against SEPT7 deficiency in early hematopoiesis, we decided to rescue the Hoxb8-immortalized cells by introducing doxycycline-inducible GFP-Sept7. We supposed that rescue of cells with a SEPT7 expressing vector construct will release mechanisms of protection from Cre-mediated recombination and will result in effective recombination of *Sept7* floxed allele (Figure 4A). Subsequently, we treated the cells with lentiviral vector particles encoding for Cre and red fluorescent protein (RFP) as a marker gene (Figure 4A). RFP/GFP double-positive Hoxb8-immortalized *Vav1-iCre*::*Sept7*^flox/flox^ HSPCs were FACS sorted and subsequently cultivated with or without doxycycline to induce or to stop GFP-Sept7 expression, respectively (Figure 4B). We did not detect any improvement of recombination and excision of the floxed *Sept7* allele in RFP/GFP double-positive Hoxb8-immortalized *Vav1-iCre*::*Sept7*^flox/flox^ HSPCs (Figure 4C). Monitoring of SEPT7 expression by western blot confirms expression of exogenous GFP-SEPT7 as well as endogenous SEPT7 even after exogenous expression of active Cre (Figure 4D). These results clearly indicate that HSPCs from *Vav1-iCre*::*Sept7*^flox/flox^ mice have developed effective mechanisms to protect at least one of the two *Sept7* floxed alleles from Cre-mediated excision. We suppose that SEPT7 is essential at the early stages of hematopoiesis, as its deletion at later stages of hematopoiesis (e.g., using *hCD2-iCre*) proves to be more effective. The resulting strong selection pressure likely drives cells to evade Cre-mediated recombination possibly via recombinase-induced DNA methylation (He and Ecker 2015; Liu et al. 2021). Interestingly, the expression of exogenous SEPT7 in Hoxb8-immortalized HSPCs reduces endogenous SEPT7 protein levels, likely due to reduced protein stability of SEPT7 monomers outside heteromeric septin complexes (Figure 4D).

**Figure 4.**
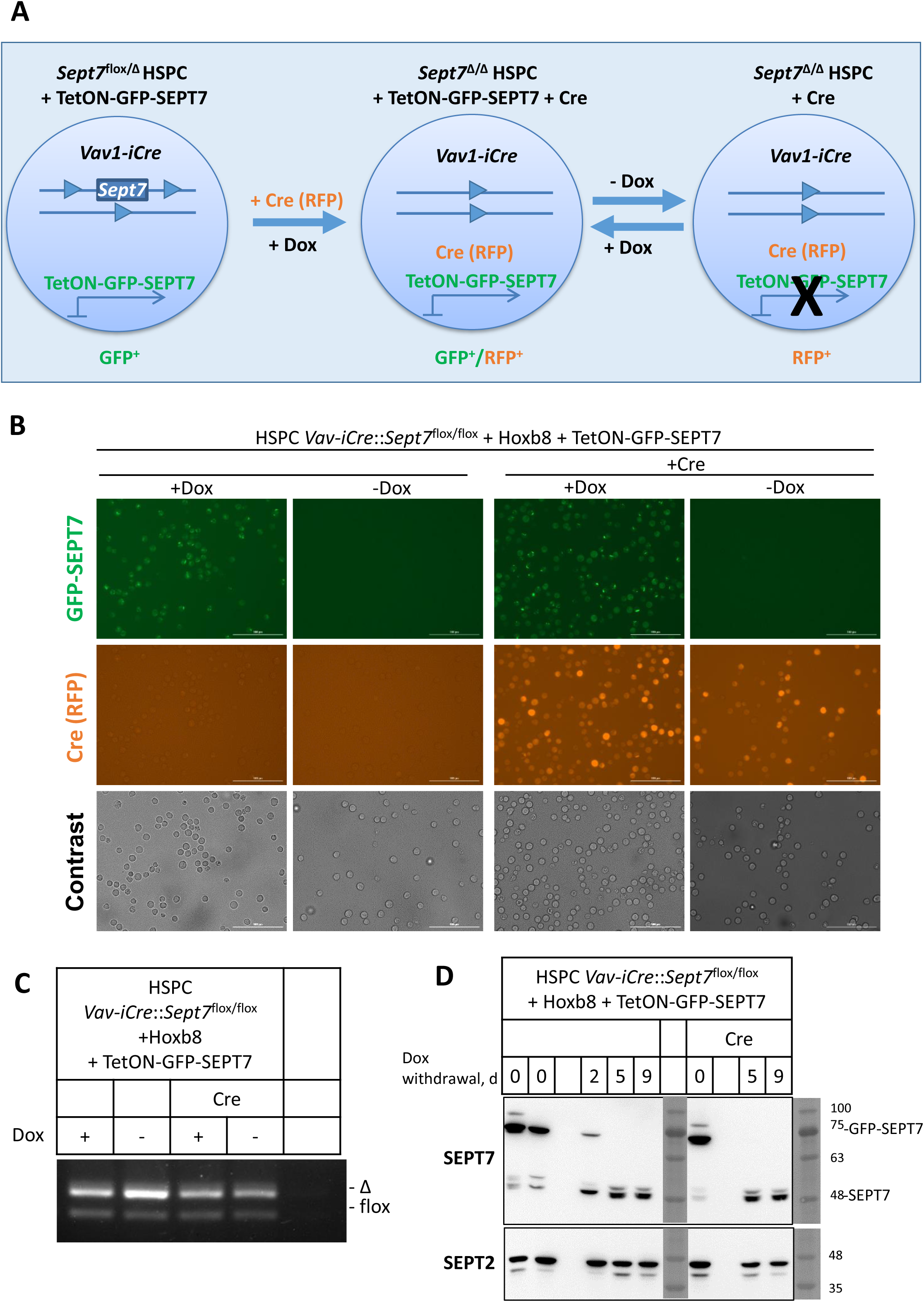
HSPCs from *Vav1-iCre*::*Sept7*^flox/flox^ mice become protected from Cre-mediated recombination. Hoxb8-immortalized HSPCs derived from *Vav1-iCre*::*Sept7*^flox/flox^ mice were double transduced first by doxycycline-inducible GFP-Sept7 and subsequently by Cre (RFP) expressing construct. Presence of doxycycline induces GFP-SEPT7 expression, while absence of doxycycline stops GFP-SEPT7 expression. GFP/RFP double-positive cells were sorted by FACS. A. Schematic representation of the GFP-SEPT7 and Cre (RFP) rescue model. B. Immuno-fluorescence analysis of single- and double-transduced HSPCs showing dox-inducible GFP-SEPT7 and RFP (Cre) signals. C. Genotyping analysis of single- and double-transduced HSPCs in the presence (+Dox) and absence (-Dox) of doxycycline indicates incomplete excision of *Sept7* floxed allele. D. Immunoblot analysis showing the doxycycline-induced GFP-SEPT7 expression in the single- or double-transduced HSPCs. Upon dox-deprivation GFP-SEPT7 disappears and endogenous SEPT7 protein levels are increased. SEPT2 was used as a loading control.

### Generation of Hoxb8 immortalized *Sept7* KO HSPCs

Since the HSPCs from *Vav1-iCre*::*Sept7*^flox/flox^ mice were protected from recombination *in vivo*, we decided to test the recombination in HSPCs from *Sept7*^flox/flox^ mice *in vitro* using the lentiviral Cre delivery as described above. To account for the possibility that SEPT7 is indispensable for HSPC proliferation, we decided to protect the cells with ectopic expression of SEPT7. The lineage-negative cells isolated from bone marrow of *Sept7*^flox/flox^ mice were first immortalized by gammaretroviral delivery of an estrogen–regulated form of Hoxb8. The Hoxb8-immortalized *Sept7*^flox/flox^ HSPCs were further transduced with a gammaretroviral vector harboring a dox-inducible GFP-SEPT7 expression cassette. These cells were then treated with lentiviral particles to deliver Cre recombinase together with RFP marker. In the double-transduced cells, Cre-expression leads to the deletion of the endogenous *Sept7* allele and these knockout cells could be specifically monitored by RFP fluorescence. We sorted RFP/GFP double-positive cells in the presence of doxycycline by FACS. Genotyping of GFP/RFP double-positive cells reveals effective recombination of *Sept7* floxed alleles (Figure 5A). Monitoring of SEPT7 expression by Western blot (Figure 5B) and by immunostaining (Figure 5C) confirmed effective depletion of SEPT7 protein. Surprisingly, we did not observe any significant difference in population doubling time between GFP-SEPT7 expressing (+Dox) and SEPT7 deficient HSPCs monitored by cell counting (Figure 5D) or by flow cytometry (Figure 5E), indicating that SEPT7 is dispensable for Hoxb8-immortalized HSPCs propagation *in vitro*. To exclude any possibility of leakage of dox-inducible GFP-SEPT7 in the absence of doxycycline, we decided to produce SEPT7-deficient HSPCs without prior reconstitution of the cells with SEPT7 expressing constructs. For that, we transduced Hoxb8-immortalized HSPCs from *Sept7*^flox/flox^ mice with a lentiviral vector expressing mCherry and Cre from a bidirectional constitutive promoter (pLBid.pA.CTE.nlsCre.minCMV.SF.mCherry.PRE) (Maetzig et al. 2010). In the transduced cells, Cre-expression leads to the deletion of the endogenous *Sept7* allele and these knockout cells could be specifically monitored by mCherry fluorescence. We sorted mCherry-positive cells expressing active Cre recombinase and confirmed efficient excision of *Sept7* floxed allele by genotyping (Figure 6A). In Cre-expressing cells SEPT7 was not detectable neither in western blot (Figure 6B) nor in immunostaining (Figure 6C) or in flow cytometry (Figure 6D). In accordance with previously published observations (Menon et al. 2014), an absence of SEPT7 was accompanied by significant reduction of SEPT2 and SEPT9 expression (Figure 6B). Population doubling time of Sept7 KO HSPCs was comparable to the original *Sept7*^flox/flox^ cells, confirming dispensability of Sept7 for *in vitro* proliferation of HSPCs (Figure 6E).

**Figure 5.**
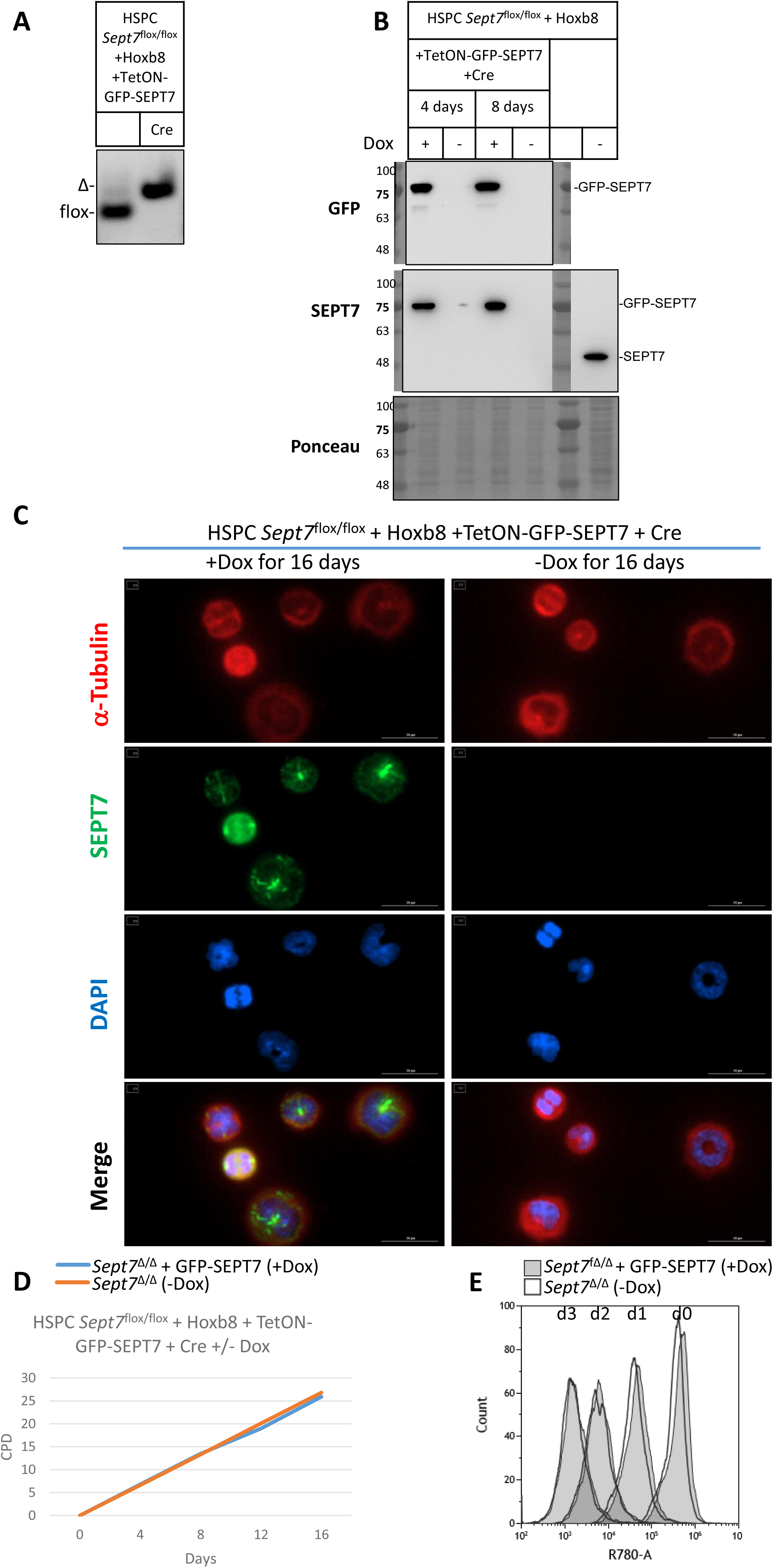
Generation of SEPT7 deficient Hoxb8 immortalized HSPCs. Hoxb8-immortalized *Sept7*^flox/flox^ HSPCs were double transduced first by doxycycline-inducible GFP-SEPT7 and subsequently by Cre (RFP) expressing construct. Presence of doxycycline induces GFP-SEPT7 expression and absence of doxycycline stops GFP-SEPT7 expression. GFP/RFP double-positive cells were sorted by FACS. A. Genotyping analysis of single- and double-transduced HSPCs in the presence of doxycycline indicates complete excision of *Sept7* floxed allele upon Cre expression. B. Immunoblot analysis showing the doxycycline-induced GFP-SEPT7 expression in the double-transduced HSPCs. Upon dox-deprivation for 4 or 8 d GFP-SEPT7 disappears and cells become negative for endogenous SEPT7, suggesting Cre-mediated excision of the floxed *Sept7* alleles. Original not transduced *Sept7*^flox/flox^ cells were used as a control. Ponceau staining was used as a loading control. C. Immuno-fluorescence analysis of double-transduced HSPC demonstrating effective depletion of SEPT7 upon dox deprivation. D. Cumulative population doublings (CPD) based on cell counting demonstrate comparable proliferation rate of SEPT7 deficient (-Dox) and GFP-SEPT7 expressing (+Dox) Hoxb8-immortalized HSPCs. E. Kinetics of proliferation of Sept7 deficient (Sept7^Δ/Δ^) and GFP-Sept7 expressing (Sept7^Δ/Δ^ + GFP-Sept7) Hoxb8-immortalized HSPCs monitored by Far Red dye dilution. Live cells were covalently labeled by Far Red Cell Tracer and each cell division resulted in half the fluorescence intensity of its parent cell. Far Red content was measured by flow cytometry at day 0, 1, 2 and 3 of cells propagation.

**Figure 6.**
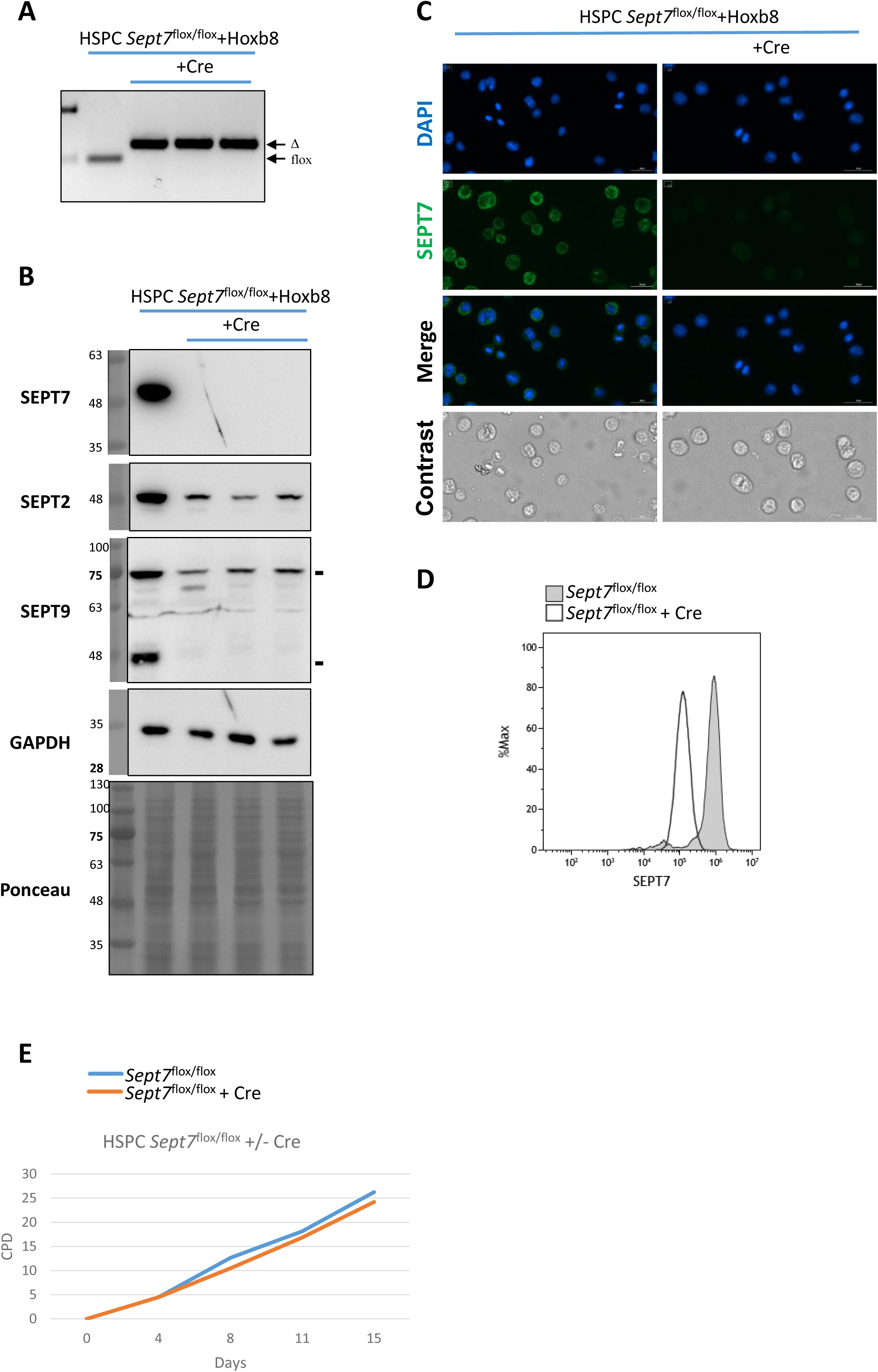
SEPT7 is dispensable for Hoxb8 immortalized HSPCs. Hoxb8-immortalized *Sept7*^flox/flox^ HSPCs were transduced with the lentiviral bidirectional construct, co-expressing Cre and mCherry and sorted for mCherry positive cells by FACS. A. Genotyping analysis indicates complete excision of *Sept7* floxed allele upon Cre expression. B. Immunoblot analysis demonstrates that Cre expression leads to effective excision of *Sept7* floxed allele. Elimination of endogenous SEPT7 is accompanied by strong reduction of SEPT2 and SEPT9 levels. GAPDH and Ponceau staining were used as a loading control. C. Immuno-fluorescent analysis confirms effective depletion of SEPT7 in *Sept7*^flox/flox^ HSPCs transduced with Cre. D. FACS analysis confirms effective depletion of SEPT7 in *Sept7*^flox/flox^ Hoxb8-immortalized HSPC stably transduced with Cre. E. Cumulative population doublings (CPD) based on cell counting demonstrate comparable proliferation rate of SEPT7 deficient (*Sept7*^flox/flox^ + Cre) and SEPT7 expressing (*Sept7*^flox/flox^) Hoxb8-immortalized HSPCs.

## Discussion

The role of SEPT7 in cytokinesis during hematopoiesis is not well understood so far (Menon and Gaestel 2015). Here, we have shown that *Vav1* driven Cre-mediated recombination is accompanied by a considerable number of *Sept7*^flox/flox^ hematopoietic cells that had not undergone recombination (escapees). These escapees are found in thymocytes, splenocytes and bone marrow and are detected at all stages of T-cell development, irrespective of higher or lower proliferative rates. In contrast, *Vav1-iCre*::*Sept7*^wt/flox^ mice demonstrated an efficiently deleted floxed allele in HSPCs and all other blood cells analyzed, indicating high efficiency of *Vav1*-driven Cre-mediated recombination. *Vav1-iCre* expresses active Cre in fetal and adult hematopoietic stem cells and all descendants (Siegemund et al. 2015). Incomplete deletion of SEPT7 from hematopoietic cells might indicate impaired repopulation activity of SEPT7 deficient cells and selective advantages of SEPT7 expressing hematopoietic cells. On the other hand, SEPT7 is efficiently eliminated at later stages of hematopoiesis, as observed in T- and B-cells from *hCD2-iCre*::*Sept7*^flox/flox^ mice, as well as in Hoxb8-immortalized hematopoietic progenitors from *Sept7*^flox/flox^ animals following transduction with a Cre-expressing construct. *hCD2-iCre* expresses active Cre at later stages of hematopoiesis if compared to *Vav1-iCre*, namely in immature and mature B and T lymphocytes (Siegemund et al. 2015). These observations suggest a requirement of SEPT7 for early stages of hematopoiesis and SEPT7 redundancy at later stages of hematopoiesis and in differentiated (mature) blood cells (Figure 7). The aging-associated changes in hematopoietic stem cells (HSCs) and hematopoiesis result from both intrinsic alterations within HSCs and extrinsic influences from the bone marrow (BM) niche (Florian et al. 2018; Mejia-Ramirez et al. 2021; Saçma et al. 2019; Guidi et al. 2017; Wang et al. 2011; Geiger et al. 2007; Kamminga and de Haan 2006). During early hematopoiesis in the fetal liver, HSCs exhibit high proliferative activity, undergoing more than a 100-fold expansion within approximately five days during embryogenesis (Ema and Nakauchi 2000; Morrison et al. 1995). In contrast, in adulthood, HSCs primarily remain in a quiescent state, which restricts self-renewal activity and helps to maintain a largely stable population of functional HSCs (Wilson et al. 2009; Cheshier et al. 1999). The distinct kinetics of HSC self-renewal across different life stages may explain our observations regarding the essential role of SEPT7 in earlier but not later stages of hematopoiesis. Polarity within HSCs is closely linked to their mode of division: young, polarized HSCs predominantly undergo asymmetric division, whereas aged, apolar HSCs tend to divide symmetrically (Florian et al. 2018). Growing evidence indicates the crucial role of septins in establishing molecular asymmetry and cell polarity in fungi and animals (Spiliotis and McMurray 2020). Thus, in budding yeast, septin barriers are dispensable for cytokinesis (Wloka et al. 2011) but are crucial for maintaining mother–daughter asymmetry, playing a fundamental role in yeast aging (Denoth-Lippuner et al. 2014; Khmelinskii et al. 2011; Gehlen et al. 2011; Shcheprova et al. 2008; Clay et al. 2014; Luedeke et al. 2005). Previous studies have shown that the Cdc42-Borg4-Septin7 axis is essential for maintaining polarity in HSCs, and that Septin7-deficient HSCs exhibit reduced engraftment potential along with characteristics of aged HSCs (Kandi et al. 2021). The data obtained in our current work confirm and expand the previously obtained data on the critical role of septin7 in HSPC function. We demonstrate that septin7 is necessary at early stages of hematopoiesis, at the time when intensive self-renewal of hematopoietic stem and progenitor cells is observed. However, at later stages of hematopoiesis, when lymphoid and myeloid precursors are predominantly proliferating, the role of SEPT7 in hematopoietic cell division becomes redundant.

**Figure 7.**
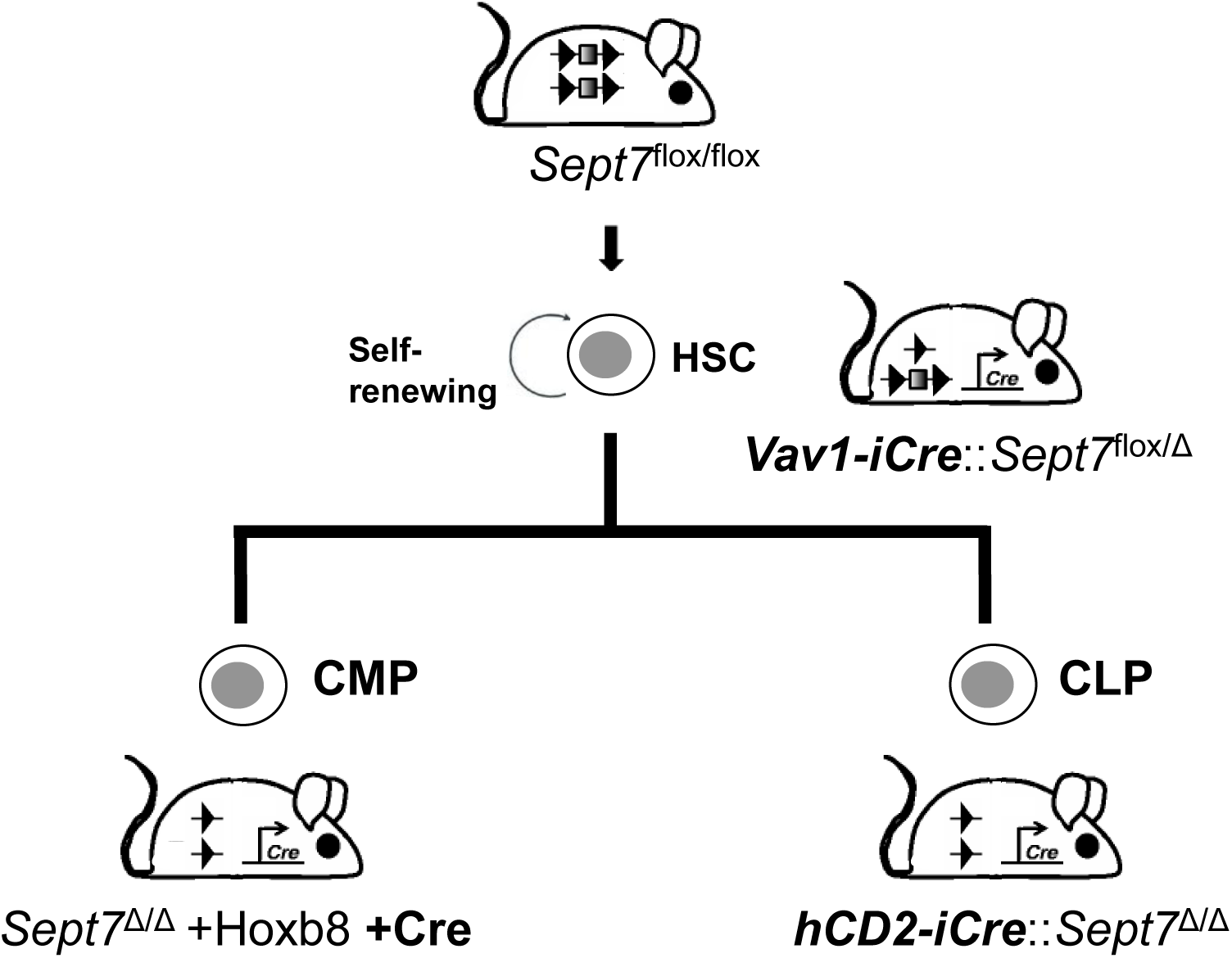
SEPT7 is indispensable at early, but not later stages of hematopoiesis. Schematic representation of the experimental results. Hematopoietic cells escape *Vav1-iCre*-mediated recombination of *Sept7*^flox/flox^ allele, indicating some stable mechanism of protection at early stages of hematopoiesis. *Sept7* is efficiently eliminated at later stages of hematopoiesis, e.g. in *hCD2-iCre*::*Sept7*^flox/flox^ T-cells or in Hoxb8-immortalized *Sept7*^flox/flox^ myeloid progenitors upon recombinant Cre expression.

## Materials and methods

### Experimental animals

All mice experiments were conducted according to German and international guidelines and were in accordance with the ethical oversight by the local government for the administrative region of Lower Saxony (permit 17/2593), Germany. *Sept7*^flox/flox^ mice (*Sept7*^tm1Mgl^) targeting the exon 4 of *Sept7* gene was reported previously (Menon et al. 2014). Lymphocyte-specific *Sept7* knockouts were generated by mating floxed animals with B6-*hCD2-iCre* mice (de Boer et al. 2003; Menon et al. 2014). To obtain hematopoietic-specific knockout of *Sept7* gene, *Sept7*^flox/flox^ mice were crossed with hematopoietic-specific *Vav1-iCre* transgene-containing mice (Kandi et al. 2021). In animal experiments sex-matched *Sept7*^wt/flox^ and *Sept7*^flox/flox^ littermates were compared. The conditional deletion of *Sept7* was tested by PCR with primer P1: 5′ GGTATAGGGGAC TTTGGGG 3′, primer P2: 5′ CTTTGCACATATGACTAAGC 3′ and primer P3: 5′ GCTTCTTTT ATGTAATCCAGG 3′ as described earlier (Menon et al. 2014).

### Antibodies and Reagents

Rabbit anti-human Septin 7, C Terminus (IBL Cat# JP18991), Polyclonal rabbit anti-septin 2 (Acris Cat# 11397-1-AP), Polyclonal rabbit anti-septin 9 (Acri, Cat# 10769-1-AP), Polyclonal goat anti-EF-2 (C-14) (Santa Cruz Biotech sc-13004-R), Monoclonal mouse anti-GFP (Santa Cruz Biotech sc-9996), Monoclonal mouse anti-GAPDH (Millipore Cat# MAB374), Monoclonal mouse anti-tubulin-α (Sigma Cat# T6199), Monoclonal anti-mouse CD4 (GK1.5) Alexa Fluor647 (BioLegend Cat# 100424, RRID:AB_389324), Monoclonal anti-mouse CD4 (GK1.5) PerCP-Cy5.5 (BioLegend Cat# 100434, RRID:AB_893324), Monoclonal anti-mouse CD8α (53-6.7) APC (BioLegend Cat# 100712, RRID:AB_312751), Monoclonal anti-mouse CD44 (IM7) PE-Cy7 (BioLegend Cat# 103030, RRID:AB_830787), Monoclonal anti-mouse CD25 (PC61) PerCP-Cy5.5 (BioLegend Cat# 102030, RRID:AB_893288), Monoclonal anti-mouse TCRβ (H57-597) BV421 (BioLegend Cat# 109230, RRID:AB_2562562), Monoclonal anti-mouse CD69 (H1.2F3) PE (BioLegend Cat# 104508, RRID:AB_313111), Donkey Anti-Rabbit IgG Alexa Fluor 488 (Invitrogen A-21206), Goat Anti-Mouse IgG Alexa Fluor 680 (Invitrogen A21057), HRP-labeled goat anti-mouse IgG (Dianova Cat# 115-035-003), HRP-labeled goat anti-rabbit IgG (Dianova Cat# 111-035-003), HRP-labeled mouse anti-goat IgG (Santa Cruz Biotech sc-2354), DAPI (Carl Roth Cat# 6335.1), Polybrene (Sigma Cat# H9268), Doxycycline (Sigma Cat# D9891), β-Estradiol (Sigma Cat# E8875), mIL-3 (Peprotech Cat# 213-13), mIL-11 (Peprotech Cat# 220-11), mSCF (Peprotech Cat# 250-03), mFlt-3L (Peprotech Cat# 250-31L).

### Cloning of gammaretroviral dox-inducible GFP-Sept7

For cloning of gammaretroviral pSERS11-T11-GFP-Sept7-PGK-M2 construct, which harbor a GFP-fused Sept7 downstream of a doxycycline-inducible promoter, we used mouse *Sept7* transcript variant 2 (NM_001205367.1) cDNA.

### Gammaretroviral virus production

The plasmids pSERS11-T11-GFP-Sept7-PGK-M2 (coding dox-inducible GFP-Sept7) or MSCV-ERHBD-Hoxb8 (Redecke et al. 2013) were co-transfected together with the ecotropic packaging vector pCL-Eco (Imgenex) into BD EcoPack 2-293 packaging cell line using PEI. Eighteen hours after transfection, the supernatant was replaced by fresh growth medium (DMEM (Gibco), supplemented with 10 % (v/v) FBS (Capricorn) and antibiotics (penicillin G (100 IU/mL) and streptomycin sulfate (100 IU/mL). Virus-containing supernatant was collected for 3-4 days every 24 h and pulled together. Supernatant was filtered through 0,45 µm filter (Sarstedt) and concentrated by ultracentrifugation in SW40-Ti rotor from Beckman Coulter: 20.000 rpm, 90 min, 4 °C. Pellet was resuspended in 2 mL IMDM (12440-053 Gibco) supplemented with 15 % FBS and antibiotics (P/S) and was filtered through 0,45 µm filter.

### pLenti-Cre virus production

The following plasmids: pLBid.pA.CTE.nlsCre.minCMV.SF.mCherry.PRE, pcDNA3.NovB2p (Sullivan and Ganem 2005), pRSV.Rev( kindly provided by Thomas J. Hope, Northwestern University, Chicago, IL), LentiGag/Pol (Schambach et al. 2006) and VSVG (Yang et al. 1995) (5 µg each) were co-transfected into 10 cm plate with 90 % confluent HEK293 cells as described elsewhere (Maetzig et al. 2010). Eighteen hours after transfection, the supernatant was replaced by fresh growth medium (DMEM (Gibco), supplemented with 10 % (v/v) FBS (Capricorn) and antibiotics (penicillin G (100 IU/mL) and streptomycin sulfate (100 IU/mL). Virus-containing supernatant was collected for 3-4 d every 24 h and pulled together. Supernatant was filtered through 0,45 µm filter (Sarstedt) and concentrated by ultracentrifugation in SW40-Ti Beckman: 25.000 rpm, 120 min, 4 °C. Pellet was resolved in 2 mL IMDM (12440-053 Gibco) supplemented with 15 % FBS and antibiotics (P/S) and was filtered through 0,45 µm filter. In some experiments commercially available pre-made lentiviral particles expressing nuclear permeant Cre recombinase under the CAG promoter and RFP-Blasticidin fusion dual marker under RSV promoter, which allows to select the positive transduced stable cells for long term culture via RFP cell sorting or via antibiotic selection (LVP577-PBS Gentarget).

### Generation and cell culture of Hoxb8 progenitor cell line

Bone marrow lineage negative cells were isolated by MagCellect™ Mouse Hematopoietic Cell Lineage Depletion Kit (R&D: # MAGM209). Cells were cultured in SFEM (StemSpam SFEM Serum-free medium Cat. 09600 STEMCELL) supplemented with P/S; 20 ng/mL IL-3; 100 ng/mL IL-11; 50 ng/mL SCF and 100 ng/mL Floxt-3L for two days prior to virus transduction. Cells were transduced on the third day: 1 mL Hoxb8 virus supplemented with 8 µg/mL Polybrene was added to 100 µL of cell suspension in 12-well plate followed by centrifugation at 2500 rpm, 30 min, 32 °C. After centrifugation 20 ng/mL IL-3; 100 ng/mL IL-11; 100 ng/mL SCF; 100 ng/mL Floxt-3L and 1 µM β-Estradiol were added to the cells. At 24 h after transduction virus was replaced with growth medium: IMDM (12440-053 Gibco) supplemented with 15% FBS; P/S; 20 ng/mL IL-3; 100 ng/mL IL-11; 100 ng/mL SCF; 100 ng/mL Floxt-3L und 1 µM β-Estradiol. Selection of transduced cells with 0,5 - 1 mg/mL Geneticin (Gibco) was started 72 h after transduction. Cells were dispensed every 3–4 d in fresh medium and transferred into new wells. Once the cell populations were stably expanding, cells were kept at concentrations between 1 × 10^4^ cells/mL medium and 1 × 10^6^ cells/mL medium.

### Western immunoblotting

Protein extracts were prepared by direct lysis of the cells with 2× Laemmli SDS sample buffer. Protein lysates were separated by SDS-PAGE on 7,5–16 % gradient gels and transferred by wet blotting to Hybond ECL nitrocellulose membranes (GE Healthcare). Western blots were blocked with 5 % powdered skim milk in PBS with 0,1 % Tween 20 for 1 h at room temperature, followed by overnight incubation with the primary antibody at 4 °C. After intensive washes with PBS containing 0,1 % Tween 20, membranes were incubated for 1 h with horseradish peroxidase-conjugated secondary antibodies at room temperature. Digital chemiluminescence images were taken by a Luminescent Image Analyser LAS-3000 (Fujifilm).

### Intracellular immunofluorescence staining

Glass coverslips were treated with 1 % Alcian blue solution for 20 min in 12-well plate and washed with PBS. Suspension cells were added in PBS and the plate was centrifuged at 2500 rpm for 10 min at 4 °C. Cells were fixed in 1 mL 4 % PFA/5 % Methanol/PBS for 15 min, washed with PBS and permeabilized with 0,25 % Triton X-100–PBS for 30 min at room temperature. Blocking was done using 4 % bovine serum albumin (BSA) for 1 h at 4 °C. Primary antibodies were used at a 1:200 dilution in 1 % BSA–PBS for 1 h at room temperature. Secondary antibodies or Alexa Fluor 647-conjugated phalloidin was used at a 1:500 dilution in 1 % BSA– PBS. Imaging was performed using Cytation1 (BioTech) with standard settings.

### Flow cytometry analysis

Thymocyte single cell suspension were generated by mashing thymi through a 100 µm sterile nylon cell strainer (Corning). The cell suspension was further filtered through a 30 µm filter, to ensure maximum removal of tissue debris. Immature thymocyte populations were characterized by staining for CD4 (Alexa Fluor 647), CD8α (APC), CD44 (PE-Cy7), CD25 (PerCP-Cy5.5) surface markers, followed by intracellular staining with TCRβ (BV421) and unconjugated anti-human Septin7 (see below). Mature thymocytes were characterized by staining for CD4 (PerCP-Cy5.5), CD8α (APC), CD69 (PE), TCRβ (BV421) followed by intracellular staining with unconjugated anti-human Septin7. For SEPT7 staining, thymocytes/splenocytes/bone marrow or hematopoietic stem cells were fixed with 3× by volume PFA (4%) at RT for 30 min. Washed and resuspended in PBS and absolute methanol was added to a final concentration of 90% by constant mixing. The methanol permeabilization was continued for 30 min on ice. After washing the cells twice with PBS, cells were resuspended in 4 % BSA-PBS and blocked at 4 °C for 30 min. Cells were stained with primary antibodies (1∶100 in 1 % BSA-PBS) at RT for 30 min. After one time washing with PBS, samples were resuspended in secondary antibody dilution (anti rabbit Alexa floxuor-488/anti mouse Alexa fluor-680 - 1∶500 diluted in 1 % BSA-PBS) and incubated for additional 30 min. Finally, cells were washed again with PBS and stained with 0,5 µg/mL DAPI for 5 minutes, resuspended in PBS, and analyzed by flow cytometry using a Beckman Coulter CytoFLEX.

### Far Red Cell Proliferation

The CellTrace Far Red kit (Thermo Fisher Cat. No. C34564) was used to monitor cell proliferation by dye dilution. 100.000 cells were washed with PBS and were resuspended in 200 µL of 1 µM FarRed solution in PBS followed by incubation for 20 min at 37 °C. Reaction was stopped by addition of 1 mL of complete Medium. After 5min incubation at room temperature, the cells were resuspended in 650 µL of growth medium and distributed 200 µL (30.000 cells) pro well in 96-well well plate. Cells were analyzed by flow cytometer with 638 nm excitation and a 780 nm emission filter using BeckmanCoulter Cytoflex. A mean Far Red peak number was quantified using Kaluza GraphPad.

### Population doubling time

2 × 10^4^ cells were repeatedly seeded in 1 mL growth medium every 3-4 d, collected and counted. Population doubling time was estimated after formula: Doubling Time = (duration * log(2))/(log(final concentration) - log(initial concentration)), where “log” is the logarithm to base n.

### Declaration of AI-assisted technologies in the writing process

During the preparation of this work the author(s) used ChatGPT and DeepL in order to improve language and readability. After using this tool/service, the author(s) reviewed and edited the content as needed and take(s) full responsibility for the content of the publication.

## Acknowledgments

We thank Dr. Matthias Ballmaier from the Flow Cytometry and Cell Sorting Resource/MHH for cell sorting. We are grateful to Prof. Hans Haecker for providing us with 3HA-ERHBD-Hoxb8-PGK-neo construct.

## Author contributions

Conceived and designed the experiments: NR, AKr and AKot. Performed the experiments: NR, NV, PG, KL, TY, AD, AKot. Analyzed the data: NR, AKr, AKot. Contributed reagents/materials/analysis tools: ASc, MGal, ASel, MA, MBM. Contributed to the writing of the manuscript: NR, AKot, AKr, MGae. Provided conceptual insight: NR, AKot, AKr, MGae.

---

**Figure S1:**
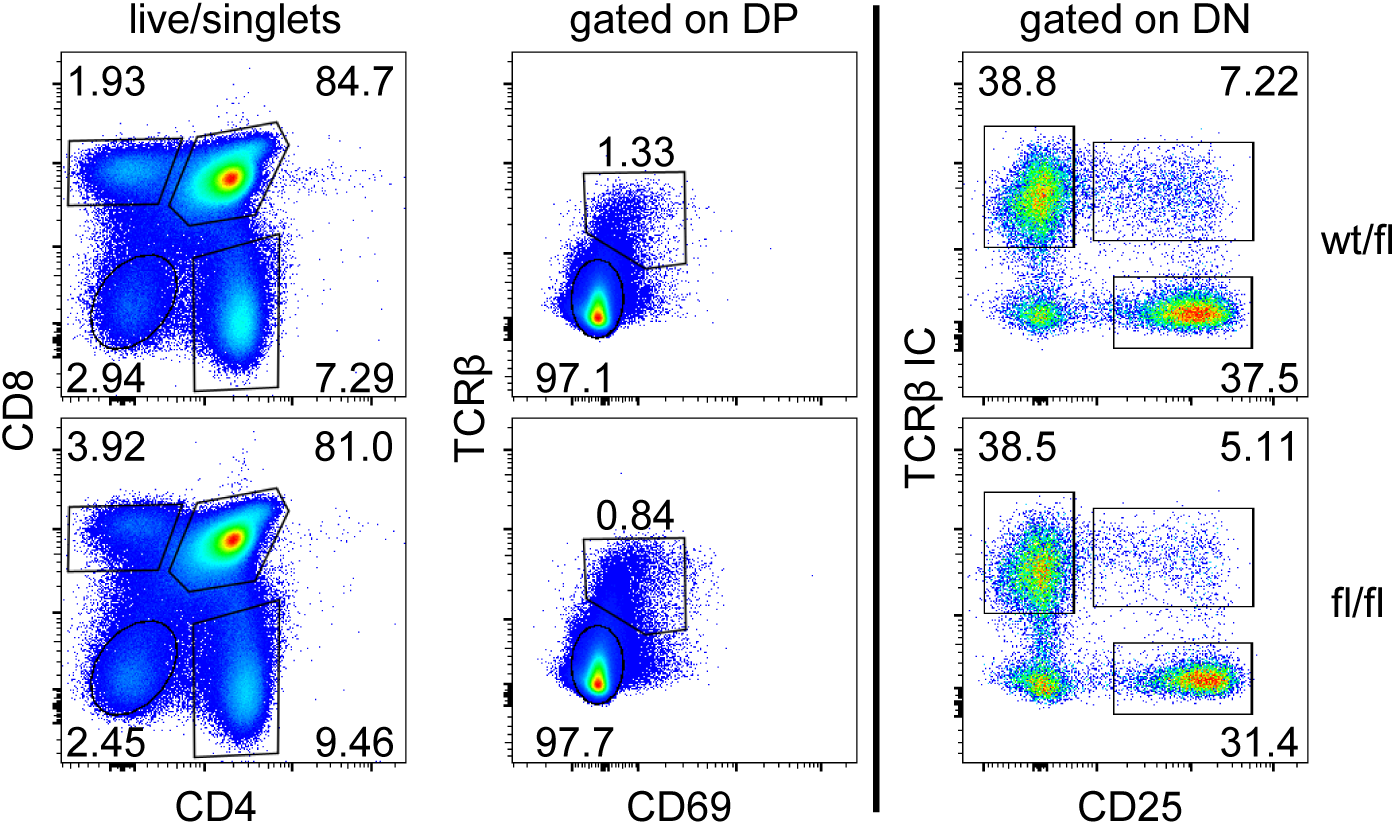
Overall development of T cells from Vav1-iCre::Sept7^flox/flox^ mice is unimpaired. Thymocytes from 12wk-old mice were analyzed for surface expression of CD4, CD8, TCR**β**, and CD69 (left and center) or CD4, CD8, and CD25 and intracellular expression of TCR**β** (TCR**β** IC, right). Populations are definfed as double-negative (DN), CD4^−^CD8^−^); double-positive (DP), CD4^+^CD8^+^; single-positive (SP), CD4^+^CD8^−^ and CD4^−^ CD8^+^; pre-selection DP, TCR**β**^−^CD69^−^; post-selection DP, TCR**β**^+^CD69^+^; DN3a, CD25^+^TCR**β**ic^−^; DN3b, CD25^+^TCR**β**ic^+^; DN4, CD25^−^TCR**β**ic^+^. Numbers adjacent to gates indicate frequencies relative to parent gates indicated on top. Data are representative of 2 12wk and 2 38wk old mice.

**Figure S2:**
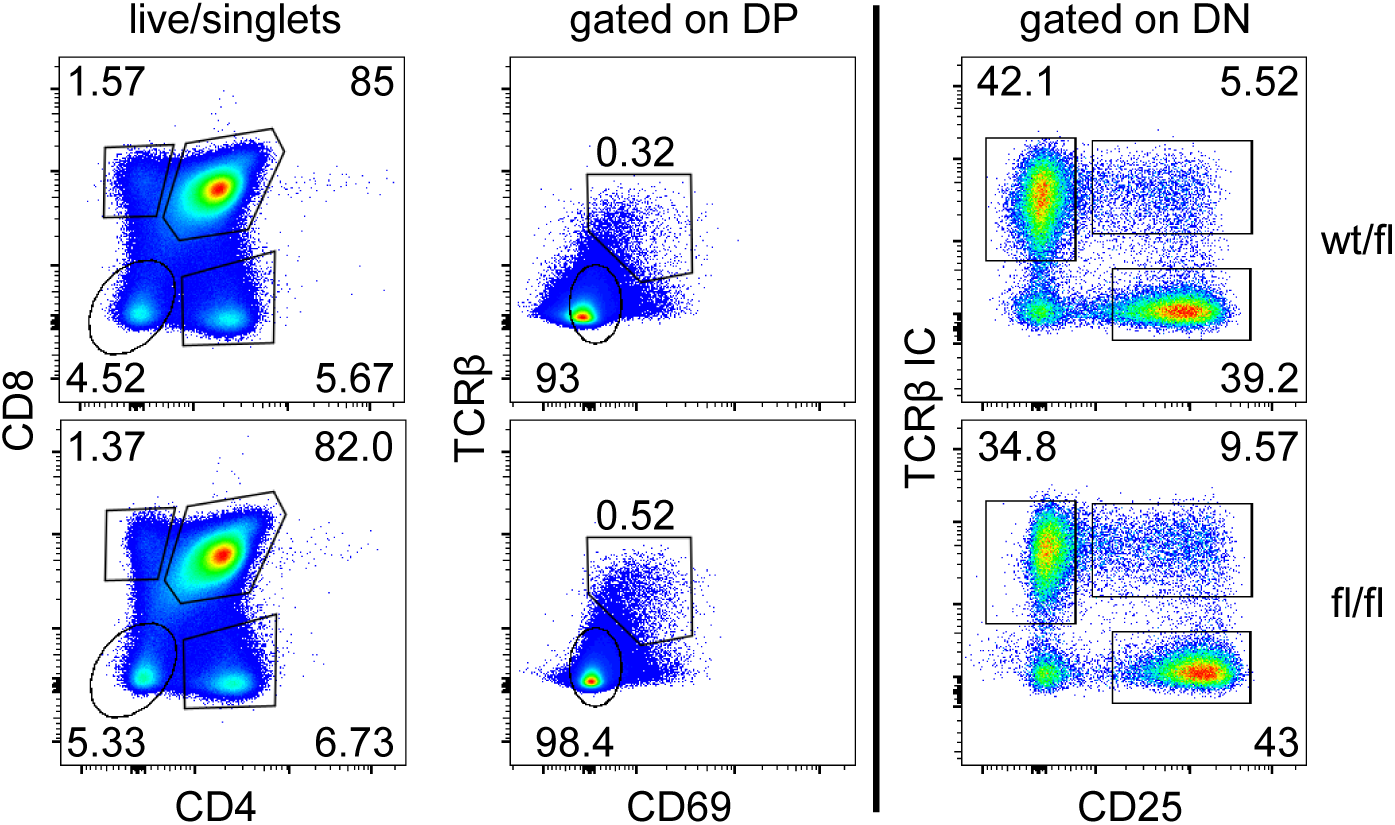
Overall development of T cells from hCD2-iCre::Sept7^flox/flox^ mice is unimpaired. Thymocytes from 14wk-old mice were analyzed for surface expression of CD4, CD8, TCR**β**, and CD69 (left and center) or CD4, CD8, and CD25 and intracellular expression of TCR**β** (TCR**β** IC, right). Populations are definfed as double-negative (DN), CD4^−^CD8^−^); double-positive (DP), CD4^+^CD8^+^; single-positive (SP), CD4^+^CD8^−^ and CD4^−^ CD8^+^; pre-selection DP, TCR**β**^−^CD69^−^; post-selection DP, TCR**β**^+^CD69^+^; DN3a, CD25^+^TCR**β**ic^−^; DN3b, CD25^+^TCR**β**ic^+^; DN4, CD25^−^TCR**β**ic^+^. Numbers adjacent to gates indicate frequencies relative to parent gates indicated on top. Data are representative of 2 14wk and 2 40wk old mice.

